# Distinct phenotype of SARS-CoV-2 Omicron BA.1 in human primary cells but no increased host range in cell lines of putative mammalian reservoir species

**DOI:** 10.1101/2022.10.04.510352

**Authors:** Manel Essaidi-Laziosi, Francisco Javier Perez Rodriguez, Catia Alvarez, Pascale Sattonnet-Roche, Giulia Torriani, Meriem Bekliz, Kenneth Adea, Matthias Lenk, Tasnim Suliman, Wolfgang Preiser, Marcel A. Müller, Christian Drosten, Laurent Kaiser, Isabella Eckerle

**Author notes:** corresponding author: Isabella Eckerle, University Hospital of Geneva, Rue Gabrielle-Perret-Gentil 4, 1205 Geneva, Switzerland.

## Abstract

SARS-CoV-2’s genetic plasticity has led to several variants of concern (VOCs). Here we studied replicative capacity for seven SARS-CoV-2 isolates (B.1, Alpha, Beta, Gamma, Delta, Zeta, and Omicron BA.1) in primary reconstituted airway epithelia (HAE) and lung-derived cell lines. Furthermore, to investigate the host range of Delta and Omicron compared to ancestral SARS-CoV-2, we assessed replication in 17 cell lines from 11 non-primate mammalian species, including bats, rodents, insectivores and carnivores. Only Omicron’s phenotype differed *in vitro*, with rapid but short replication and efficient production of infectious virus in nasal HAEs, in contrast to other VOCs, but not in lung cell lines. No increased infection efficiency for other species was observed, but Delta and Omicron infection efficiency was increased in A549 cells. Notably replication in A549 and Calu3 cells was lower than in nasal HAE. Our results suggest better adaptation of VOCs towards humans, without an extended host range.

## Introduction

Severe Acute Respiratory Syndrome Coronavirus 2 (SARS-CoV-2) is the etiological agent of coronavirus disease 19 (COVID19) that has caused one of the biggest public health challenges of modern times (1). Few mutational changes were observed in SARS-CoV-2 during the first year of the pandemic, most notably the Spike D614G mutation. This mutation enhanced angiotensin converting enzyme-2 (ACE-2) binding, providing a fitness advantage, and was responsible for the first pandemic wave (2, 3). By late 2020, SARS-CoV-2 variants had emerged, characterised by numerous mutations, mainly in Spike. These were classified as variants of concern (VOC), variants of interest (VOI) or variants under monitoring (VUMs), based on their genetic, clinical and epidemiological characteristics (4). To date, five VOCs have been designated: Alpha, Beta, Gamma, Delta and Omicron. VOCs are characterized by a rapid increase in case numbers, quickly outcompeting earlier strains in their region of emergence (5-9). Omicron has the most observed mutations, with the majority located in Spike, causing the strongest escape from prior immunity so far (10). It also shows signs of higher transmissibility and secondary attack rates (11). In addition, VUMs and VOIs were co-circulating along with VOCs, including the (former) VOI Zeta used in our study here, that arose alongside the Gamma VOC in South America during a surge in local cases, but nearly all have disappeared (12).

Variant evolution does not yet follow a pattern, and most variants have not evolved from each other, but arose independently from basal circulating viruses (13). Indeed, Omicron is most closely related to a virus that was circulating in mid-2020. There are three main hypotheses on VOC origins (14). The first holds that they developed in populations that were not covered by genomic surveillance. The second involves reverse zoonotic events, with transmission from humans into intermediate hosts, and then back to humans. While a large range of animal species are known to be susceptible to SARS-CoV-2, no plausible VOC progenitors have been found in animals (15). The third hypothesis describes intra-host evolution during chronic infections in immunocompromised patients. Such cases have already been reported, with prolonged shedding of mutated infectious virus (16-18).

VOCs have shown distinct epidemiological and partial clinical differences. While genomic surveillance can inform about mutations and the phylogenetic relationships, it cannot directly conclude on biological properties resulting from these mutations, thus the *in vitro* assessment of phenotypic differences is of immediate relevance whenever new variants arise. Understanding the mechanisms of enhanced fitness, the origins of VOCs and their risk for reverse zoonotic events are crucial for the further control of this pandemic. Here, we have assessed replicative capacity and potential *in vitro* phenotypes of ancestral SARS-CoV-2 (B.1), VOCs Alpha, Beta, Gamma, Delta, Omicron (BA.1) and former VOI Zeta on human primary cells and lung-derived cell lines and on mammalian cell lines derived from bats, rodents, insectivores and carnivores to investigate potential host range.

## Results

### Comparative replicative capacity of SARS-CoV-2 variants in human cell culture models of the respiratory tract

To compare viral phenotypes, we infected primary human airway epithelial cells (HAE), derived from the nasal epithelium and differentiated *in vitro* in 3-dimensional (3D) air-liquid interface cultures and monolayer (2D) cultures with SARS-CoV-2 clinical isolates at the multiplicity of infection (MOI) of 0.1 at 37°C and 33°C (**Figure 1**). Upon infection of HAE cultures and incubation at 37°C, we observed a rapid increase at 24h post infection for Omicron, from mean log_10_ RNA copy numbers/mL (RNAc/mL) of 7.8 to a peak of 11.8, while all other isolates reached similar peak values at around 96 hours. Omicron, in contrast, showed a reduction of viral RNAc/mL at 96h to 10.8 log_10_ RNAc/mL. No differences in replication kinetics were observed between B.1, Alpha, Beta, Gamma, Delta and Zeta, in all of which log10 RNAc gradually increased over the first 4 days.

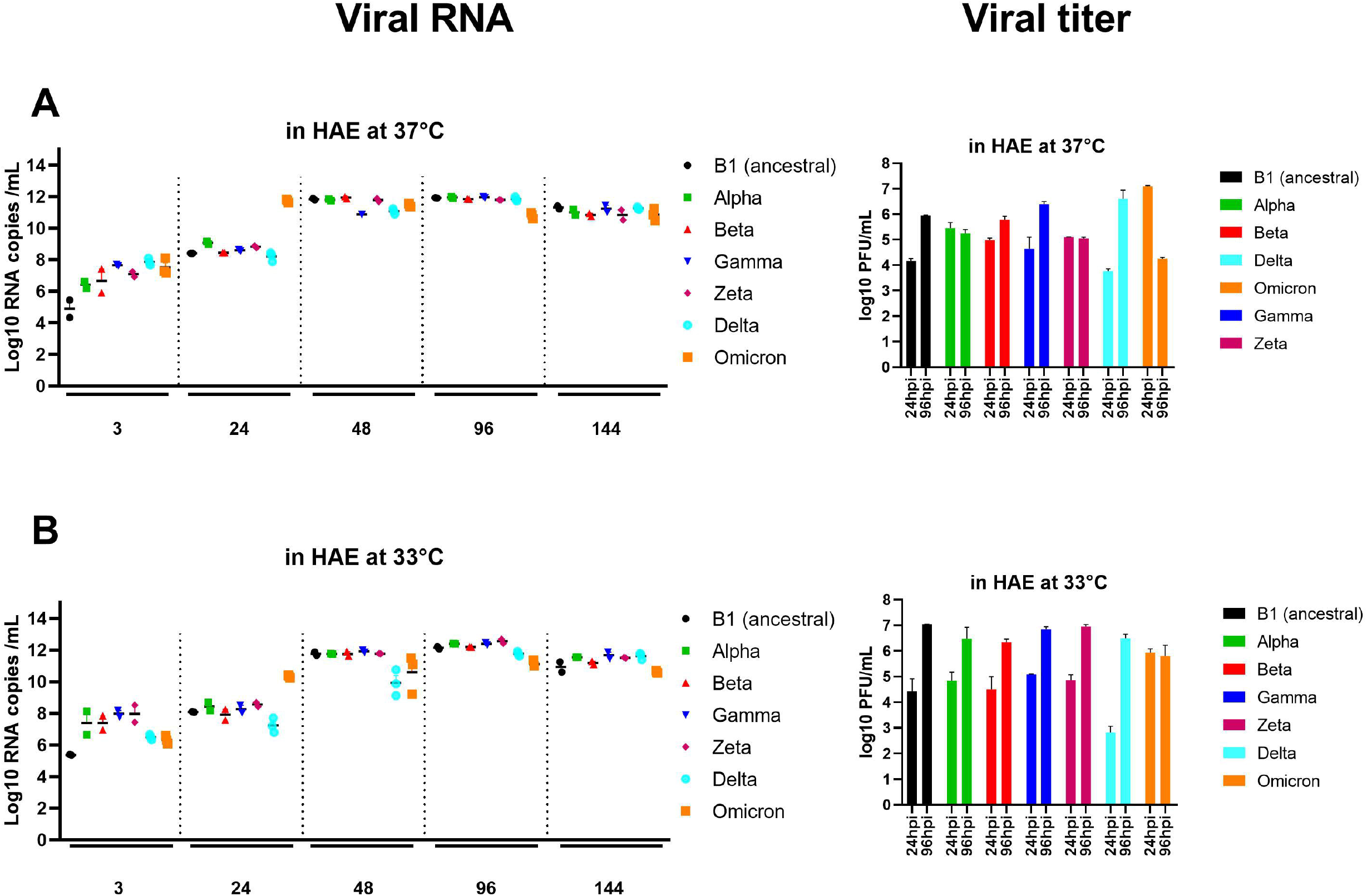

When assessing infectious viral particles by plaque-forming assay at 37°C, Omicron reached peak infectious viral titer of 7.1 log_10_ plaque-forming units (PFU) after 24h with a rapid decline to only 4.3 after 96h. On the contrary, B.1, Beta, Gamma and Delta showed an increase in PFU from 24 to 96 h, but peak PFU titers stayed below that of Omicron. The Alpha and the Zeta variants did not show strong increases, with PFUs for 24 and 96h staying constant and below those of the other variants. The strongest increase was observed for Delta, which showed the lowest infectious titer at 24h of 3.8, which increased to 6.6 mean log_10_ PFU/mL.

At 33°C incubation temperature, Omicron also showed a fold increase of 4.0 mean log_10_ RNAc/mL at 24h, but RNA levels increased further with a peak at 96h of 11.1 mean log_10_ RNAc/mL. Similarly, the other variants including B.1 reached their peak RNA levels at 96h, but with higher RNA levels than that of Omicron, ranging between 11.8 and 12.6 mean log_10_ RNAc/mL. Relative to 37°C, at 33°C viral RNAc/mL were 0.3-0.8 log_10_ higher for B.1, Alpha, Beta, Gamma and Zeta, but not for Delta and Omicron, which both showed slightly lower viral loads at 33°C. Similarly, PFUs were higher for B.1, Alpha, Beta, Gamma and Zeta at 33°C, but similar for Delta and lower for Omicron. No higher PFU titer was observed for Omicron at 24h, and similar infectious titers were found for 24 and 96 hpi (5.8-5.9 mean log10 PFU/mL).

The same experiments were performed in Calu3 cells derived from a human lung adenocarcinoma (**Figure 2**). Replication kinetics at 37°C were similar for all viruses, including Omicron, with peak RNA levels reached at 48hpi. Peak RNA levels were comparable between B.1, Alpha, Beta, Gamma and Zeta, ranging from 10.7 to 11.3, but higher for Delta and Omicron (12.1 for both). No early peak for Omicron, such as in HAE cultures, was observed. In contrast to HAE cultures, PFU titers peaked rapidly in this cell line at 37°C, with higher infectious viral loads at 24h post infection than at 96hpi. Only Omicron did not show an early PFU titer peak, but instead showed an increase from 24 to 96hpi, with peak PFU titers remaining lower than those of the other variants (mean 5.1 log_10_ PFU/mL).

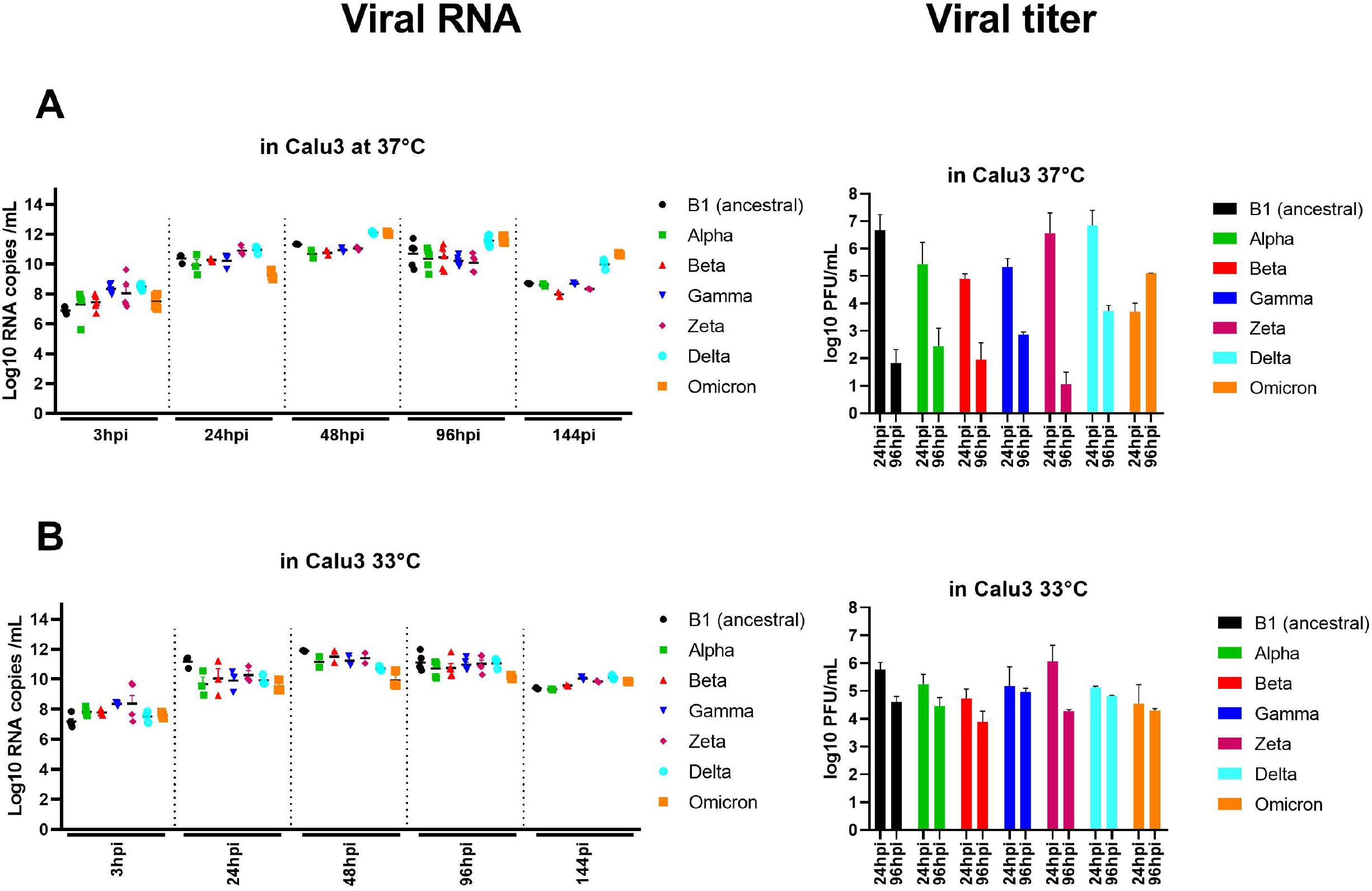

Viral replication at 33°C showed a homogeneous replication pattern between variants, with peak RNA levels for B.1, Alpha, Beta, Gamma and Zeta reached at 48hpi in the range of 11.1-11.9, while peak RNA levels for Delta and Omicron were delayed with a peak 96hpi with titers of 11.0-10.1, respectively. Among all variants, Omicron showed the lowest replication with ca. 1 log_10_ RNA copies/mL lower than Delta and up to 2 log_10_ RNA copies/mL lower compared to B.1, Alpha, Beta, Gamma and Zeta.

At 33°C compared to 37°C, B.1, Alpha, Beta, Gamma and Zeta had higher peak RNA levels by 0.3 to 0.6 log_10_ RNAc/mL, while Delta and Omicron had 1.0 and 2.0 log_10_ RNAc/mL lower peaks, respectively. PFU titers were lower in Calu3 at 33°C, but infectious viral titers 24hpi were still higher than at 96hpi.

#### Replicative capacity of B.1, Delta and Omicron in mammalian cell lines

In order to assess the replicative capacity of B.1, Delta, and Omicron BA.1, we infected 17 non-human mammalian cell lines, from 11 wild or domestic species (**Table 1**). Species include a range of European wildlife. Cells were inoculated at a MOI of 0.1 and 37°C. The supernatant’s viral RNA was quantified after 4 days, along with the PFU titer if viral RNA had increased. Human A549 (with low expression of ACE-2) cell lines and Vero-E6 cell lines (primate) were infected in parallel as controls (**Figure 3**). Efficient replication was observed for all three viruses in a rabbit kidney cell line (RK-13), with RNA levels between 10.3-11.5 log_10_ RNAc/mL at 4 dpi. Upon titration, infectious titers were between 5.8-7.0 PFU/mL.

**Table 1:** List of cell lines used in the study. *confirmed by the sequencing of the cytochrome C subunit oxidase I gene.

| Designation | Species of Origin | Species name (latin)* | Mammalian order | Geographic range | Organ of Origin |
| --- | --- | --- | --- | --- | --- |
| **RF Kidney** | Greater horseshoe bat | *Rhinolophus ferrumequinum* | *Chiroptera* | Europe | Kidney |
| **RE Kidney** | Mediterranean horseshoe bat | *Rhinolophus euryale* | *Chiroptera* | Europe | Kidney |
| **RE Trachea** | Mediterranean horseshoe bat | *Rhinolophus euryale* | *Chiroptera* | Europe | Trachea |
| **RCL.B Trachea** | Geoffroy's horseshoe bat | *Rhinolophus clivosus* | *Chiroptera* | Africa | Trachea |
| **Pip Trachea** | Common pipistrelle | *Pipistrellus pipistrellus* | *Chiroptera* | Europe | Trachea |
| **NE-R** | American mink | *Mustela vison* | *Carnivora* | America, introduced in other areas of the world | Foetus |
| **MV1-Lu** | American mink | *Mustela vison* | *Carnivora* | America, introduced in other areas of the world | Lung |
| **FUFE-R Fbr** | Red fox | *Vulpes vulpes* | *Carnivora* | Europe | Embryo |
| **Crocidura** | Lesser white-toothed shrew | *Crocidura suaveolens* | *Soricomorpha* | Europe | Trachea |
| **Igel 2 REC.B** | European hedgehog | *Erinaceus europaeus* | *Erinaceomorpha* | Europe | Kidney |
| **Mygla AEC.B** | Bank vole | *Myodes glareolus* | *Rodentia* | Europe | Trachea |
| **MPFLU-1-R** | Domestic ferret | *Mustela putorius furo* | *Carnivora* | Europe | Lung |
| **MPFN-1-R** | Domestic ferret | *Mustela putorius furo* | *Carnivora* | Europe | Kidney |
| **KAN-2-R** | Rabbit | *Oryctolagus cuniculus* | *Lagomorpha* | Europe | Lung |
| **RK-13** | Rabbit | *Oryctolagus cuniculus* | *Lagomorpha* | Europe | Kidney |
| **ZP** | Rabbit | *Oryctolagus cuniculus* | *Lagomorpha* | Europe | Embryo |
| **R9ab** | Rabbit | *Oryctolagus cuniculus* | *Lagomorpha* | Europe | Kidney |
| **A549** | Human | *Homo sapiens* | *Primates* | Global | Lung |
| **VERO-E6** | African green monkey | *Chlorocebus aethiops* | *Primates* | Africa | Kidney |

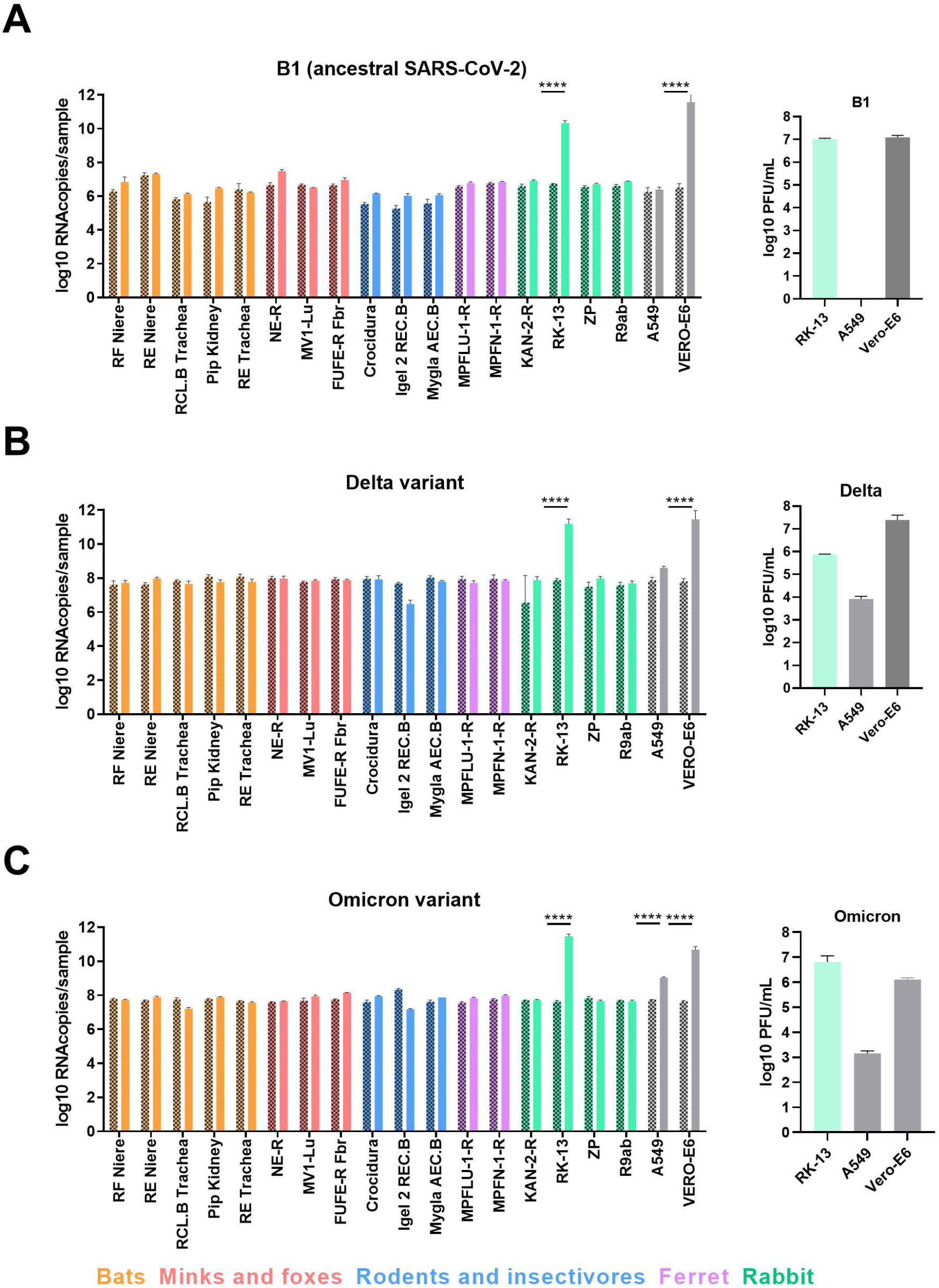

Vero-E6 cells, which were infected in parallel, showed an increase in viral RNA between 10.7-11.6 log_10_ RNA copies/mL, resulting in 6.1-7.0 PFU/mL. No other non-human mammalian cell lines from the species of bats, rodents, minks and foxes showed increased RNA indicating efficient replication, further confirmed by a lack of increase in sub-genomic RNA for all three viruses (**Figure S1**). In contrast, human A549 cells did not show an increase of genomic nor by sub-genomic viral RNA upon infection with B.1, but were efficiently infected by Delta and Omicron with respective increases in log_10_ RNA copies/mL at 4 dpi of 0.7 and 1.3 and infectious virus titers of 3.9 and 3.2 PFU/mL.

#### Comparison of ACE-2 orthologs from animal species

In order to understand differences between ACE-2 receptor binding residues of tested cell lines, we decided to compare ACE-2 amino-acid sequences between the studied species (**Table 2**). These sequences were either available in GenBank/NCBI or obtained by sequencing of the cell lines used when no sequence was available yet (see material and method). The alignment and comparison of key ACE-2 orthologue residues involved in binding to the spike protein showed no difference between *Homo sapiens* and *Chlorocebus aethiops* (Vero-E6). Of the other mammalian species, rabbits and Red foxes showed the highest (76.92%) homology to human ACE-2. Bats, shrews and hedgehogs showed the lowest homology rates (50-65.3%). Amino-acids F28, E37, L45, N53, N330, K353, D355, R357 and R393 were highly conserved between species while amino-acids at positions 38, 41, 83, 27, 322, 329 and 354 were rather variable.

**Table 2:**
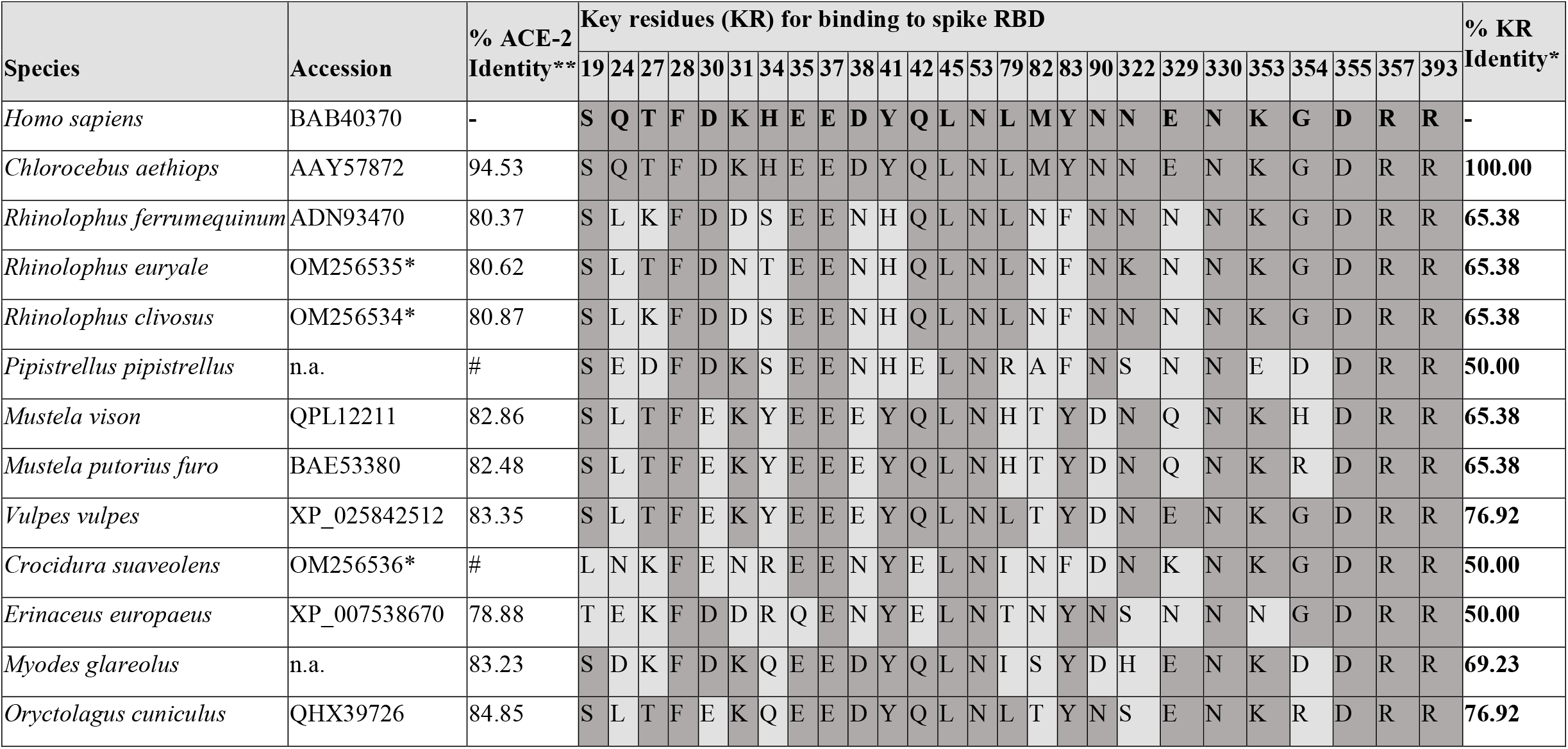
Comparison of ACE-2 key residues for SARS-CoV-2 binding in animal cells. Full length and key residues of ACE-2 protein sequences were compared between the different animal species including human. Eight sequences were already available. Sequences from *Rhinolophus euryale, Rhinolophus clivosus* and *Crocidura suaveolens* were newly added in this work (cf. accession codes). *sequence obtained in this study **percentage of identity of ACE-2 full sequence or key residues (KR) *versus H. sapiens*. #: partial sequences. n.a.: no accession number available. Differences compared to the human sequence are highlighted in light grey.

## Discussion

Since its emergence in 2019, SARS-CoV-2 has been in constant evolution, frequently giving rise to new variants (19). The emergence of several VOCs, including the latest Omicron has stressed the need for continuous surveillance and rapid phenotypic assessment to guide public health measures. A plethora of studies compared SARS-CoV-2 variants either by live virus or pseudo-viral assays using human *in vitro* models such as cell lines or organoids, or animal models, with the aim to provide rapid answers on the altered biological properties of novel variants (11, 20-22). Differentiated 3D tissues of the human respiratory tract and organoids are considered as one of the most relevant models for such studies as they are human-derived, readily available and most closely recapitulate the *in vivo* situation (23). Nevertheless, widely available immortalized cells are commonly used to study virus tropism and pathogenicity and decipher the mechanisms involved, and can reflect selected aspects. A range of immortalized cell lines of wildlife species have become available in recent years, allowing study of the susceptibility of different hosts, especially those that are not experimental animal models (24-27). In our study, we extensively compare the replication of isolates of seven SARS-CoV-2 lineages in cell culture models of the human respiratory tract and in cell lines obtained from domestic and wildlife species, with a focus on European species.

In nasal HAE cells, our findings clearly demonstrate a distinct BA.1 phenotype compared to the other variants, with shorter but faster replication to higher levels, and early efficient production of infectious virus. The early and efficient replication, in addition to its immune evasion properties, could contribute to Omicron’s contagiousness and high secondary attack rate (28, 29). However, in a recent clinical study, we did not observe more infectious viral shedding for Omicron at symptom onset compared to Delta (30). These differences could be reflected by the fact that our HAE model allowed studying the very early phase of virus replication that is undetected by diagnostic testing, as clinical samples are usually collected after symptom onset. Furthermore, adaptive immunity is lacking in our HAE model, which may mitigate replication, especially given the infection rates in vaccinated or previously infected individuals. Although Omicron quickly reaches high viral levels, both RNA levels and PFU titer rapidly declines in HAE cells, and are lower than levels for Delta at 96 hours, in agreement with clinical observations.

Agreeing with our findings, early and rapid replication of BA.1 was also seen in human *ex vivo* bronchus but less in lung parenchyma (21). Less efficient cleavage of the Omicron spike was suggested to reduce fusogenicity (31). In our study, we have also seen less efficient replication in lung-derived cells versus HAE cultures, although there are limits to their comparisons. The less efficient replication in the lower respiratory tract might explain the reduced clinical severity of BA.1 (22, 31), while the early efficient replication may contribute to efficient community spread. Additionally, its efficient replication could be explained by an enhanced cell host entry thanks to improved ACE-2 binding and more efficient endocytosis, rather than lower sensitivity to interferon responses (32).

Better replication was shown for ancestral SARS-CoV-2 at 33°C in nasal and lung *in vitro* models (33), and we also observed more efficient infectious virus production at lower temperature for Alpha, Beta and Zeta, but not for Delta and Omicron (Figure1). Similarly, no replication differences between 33°C and 37°C were observed for Omicron in organoids of bronchi and lung in another study (21).

We also investigated host tropism in cell culture models. SARS-CoV-2 has already demonstrated high promiscuity by infecting multiple animal species, with both animal-human spill-backs and establishment of new animal reservoirs (15, 34-36). In general, inter-species transmission is a well-known feature of coronaviruses, considerably contributing to their pandemic potential, and in the case of SARS-CoV-2 facilitated by conservation of the host receptor ACE-2 across mammalian species (37-39).

As with any new emerging virus, establishment of new host reservoirs poses a risk for further mutation of the virus, complicates control of transmission and eventually leads to spill-backs into humans. Both an origin as well as an increased risk for spill over into new reservoirs of new variants has been discussed (40). We did not see an increased host range for Delta or BA.1 in our animal cell lines, which included a range of European small mammals and several species of bats of the family *Rhinolophidae. Rhinolophidae* are known to host diverse SARS-related coronaviruses, and the closest known relative of SARS-CoV-2 was found in an Asian species (41, 42). In contrast, we investigated two European species, *R. ferrumequinum* and *R. euryale*, and a South African species, *R. clivosus*, with no signs of SARS-CoV-2 replication in kidney and tracheal cell lines from these bats. Nor did the cells of a more ubiquitous non*-Rhinolophidae* bat, *P. pipistrellus*, show any signs of replication by B.1, Delta or Omicron. Other studies have shown SARS-CoV-2 does not readily replicate in a range of bat cell lines including *Rhinolophus* cells (43) while one study found signs of replication in a kidney cell line from an Asian Rhinolophus species (44). Another study found bat cell lines to be only weakly susceptible, which could be overcome by expression of hACE-2, similar to what was observed for SARS-CoV (45, 46).

The only susceptible cell line from our panel was derived from a rabbit kidney, which replicated equally well in B.1, Delta and Omicron, but no other animal cell lines showed signs of efficient replication. Indeed, rabbits have been used as susceptible experimental animal models (47). However, three other rabbit cell lines, did not show signs of replication. Surprisingly, we were not able to reproduce efficient infection of cells originating from minks and ferrets, although SARS-CoV-2 was reported to infect both species and has caused outbreaks in minks farmed for fur (48).

Other studies have assessed the host range of SARS-CoV-2 mostly using ACE2-transgenic cell lines and pseudotyped viruses, but fewer studies have investigated VOCs, and results between studies were conflicting (49). A study using lentiviral pseudotypes and transgenic animal ACE2-expressing cell lines suggested Omicron’s tropism extended into domestic avian species, *Rhinolophus* bats and mice (50). A study using a similar approach suggested broad entry of ancestral SARS-CoV-2 into a range of animal species (cat, dog, cow, horse, camel, hamster, rabbit but not mink) as well as enhanced entry of Alpha and Beta variants (51). One study investigated well-differentiated airway cultures from a range of animals (some of which were also studied here) and found ancestral SARS-CoV-2 replication in cells derived from rhesus macaques and cats, but not in those derived from ferrets, dogs, rabbits, pigs, cattle, goats, camels, llamas and two neotropical bat species (52).

Discrepancies between *in vitro* findings and naturally observed infections or animal experiments could be because: the culture models do not accurately reflect the site of replication *in vivo*, receptor expression is reduced in cell culture, differences in the required infection dose, body temperature, or SARS-CoV-2 uses different receptors in some animal species. Results obtained with pseudotyped viruses and transgenic ACE-2 expression could furthermore differ from wild-type virus assessment in unaltered cell lines.

In addition to infection experiments, several modelling and bioinformatic approaches have tried to identify susceptible species by comparing ACE-2 sequences (37, 53). The *in-silico* prediction of ACE-2 affinity with spike’s RBD previously identified of more than 500 animal species as potentially susceptible to SARS-CoV-2 (54). Others have found that the predictive power of ACE-2 sequences is rather low and can be misleading due to biased ACE-2 sequence availability (55). Therefore, phenotypic assessment in cell lines, although not perfectly reflecting the *in vivo* susceptibility, can complement such bioinformatic studies. Cell lines from wildlife species such as bats can provide both biological infection data and a source for ACE-2 sequencing, as done here.

Despite its important variability across species, ACE-2 appears more genetically stable compared to TMPRSS2, a type II transmembrane serine protease involved in virus-cell fusion during virus entry, which had shown less identities between human and animals with possible partial-to-total TMPRSS2 gene loss in some vertebrates (56).

Together with data from animal cells, our observations support a human origin of Omicron rather than a reverse zoonosis. Although bats are considered to be the original hosts, SARS-CoV-2 does not readily replicate in bats cells because of the divergent RBD sequences compared to close relatives in bats, like RaTG13.

In conclusion, using cell models, we showed that Omicron has the strongest phenotypic differences compared to the previous variants, and likely evolved in humans rather than animals. Such cell culture models can help better understand SARS-CoV-2 infections, including VOCs, in humans. We also showed the relevance of cell models from a variety of species, even if not perfectly reflecting *in vivo* situation, to assessing the risk of zoonotic spillback.

## Material and methods

### Human and animal Cell lines

Commercially available nasal HAE, called “MucilAir™”, were purchased from Epithelix SARL [www.epithelix.com]. They are maintained in air-liquid interface culture where the medium (MucilAir™ medium, Epithelix) is supplemented by the basal chamber and the apical surface of the tissue is in contact with air (see (57) for details).

All used animal cell lines are summarized in Table1. The corresponding species were confirmed by sequencing of the cytochrome C subunit oxidase I gene (58, 59). Five additional standard cell lines were used in this study as controls or for viral stock production, including Calu3 (human lung cancer cell), Vero-E6 (African Green Monkey kidney), A549 (human lung carcinoma), A549 cells overexpressing ACE-2 receptor and Vero-E6 overexpressing TMPRSS2. All immortalized cells were cultured in monolayer using medium (MEM Glutamax, Gibco) supplemented with 10% (cell maintenance) or 2% (infections) Fetal Bovin Serum (FBS, Gibco), 1X penicillin-streptomycin (Gibco) and 1% non-essential amino-acids (MEM NEAA, Gibco). All cell cultures were performed at 37 °C under 5% CO_2_.

### Viruses

SARS-CoV-2 isolates were obtained from clinical samples collected in the outpatient testing Centre of the University Hospitals of Geneva as described previously (29). All SARS-CoV-2 isolates used are summarized in Table S1. B.1 and the variants Alpha, Gamma, Zeta and Delta were isolated after one passage in Vero-E6 cells. One additional passage allowed viral stock production in the same line. Vero-E6 was less permissive to Beta and Omicron variants. Hence, Beta was isolated in A549-ACE-2, after a second passage in mixed Vero-E6:A549-ACE-2 (1:1) cells, the viral stock was produced in Vero-E6 cells. Omicron was isolated after 2 passages in Vero-TMPRSS2. All viral stocks were titrated in Vero-E6 and sequenced.

### Infections of animal and human cells

Infection assays were performed at 37°C or 33°C at 5% CO_2_. HAE reconstituted *in vitro* (3D culture) or immortalised cell lines in 2D culture, including Calu3 and animal cell lines (Table 1), were tested for infections as previously described (20) at 37°C and an MOI of 0.1 using seven SARS-CoV-2 isolates (Table S1). Cells were washed after 1h (Calu3 and animal cells) or 3h (HAE). Supernatant (Calu3) or apical washes (HAE) were collected at different times after infection. Infected animal cells were lysed using NucliSens easyMAG lysis buffer (BioMérieux) at 4 dpi.

### Quantification of viral RNA

In order to determine the viral load from collected samples, RNA was extracted with NucliSens easyMAG (BioMérieux) and quantified by quantitative real time PCR (RT-qPCR) using SuperScript™ III Platinum™ One-Step qRT-PCR Kit (Invitrogen) in a CFX96 Thermal Cycler (BIORAD). Real time RT-qPCR was performed using specific sets of primers and probes targeting either genomic or sub-genomic viral RNA as previously described (60, 61)

### Assessment of infectious titer

Infectious titer of collected samples was assessed by plaque assays performed at 37°C and 5% CO_2_ as previously described (20). Briefly, Vero-E6 seeded in monolayer of 2×10^5^ cells/mL in 24-well plates were inoculated 2h after using serially diluted samples. The inoculum was removed and replaced by fresh medium (DMEM 10%FBS, 2mM L-glutamine, 1% penicillin-streptomycin all from Gibco) 1:1 mixed with 2.4% Avicel (RC-581, FMC biopolymer) one hour later. Cells were fixed using 6% paraformaldehyde (Sigma) 24h later at least 1h at RT and stained with crystal violet (Sigma). PFU were counted from each dilution in order to determine the infectious titer (in PFU/mL) for each sample.

### Comparison of ACE-2 orthologues’ protein sequences

For the species *Homo sapiens, Chlorocebus aethiops, Rhinolophus ferrumequinum, Mustela viso*n, *Mustela putorius furo, Vulpes vulpes, Erinaceus europaeus* and *Oryctolagus cuniculus*, ACE-2 amino-acid sequences were available in NCBI. *Pipistrellus pipistrellus* ACE-2 protein sequence has been kindly shared by Joanna Damas and Harris A Lewin (37), obtained from an original sequence from the Zoonomia Consortium. For the species *Crocidura suaveolens, Rhinolophus clivosus* and *Rhinolophus euryale* the protein sequences obtained in this study were predicted from ACE-2 nucleic sequences after RNA trizol-extraction from cell lysates and amplification by RT-PCR using primers (Microsynth) listed in Table S2. The protein sequence for *Myodes glareolus*, was predicted from a reference transcriptome (62). Contrarily to other species, ACE-2 from *Crocidura suaveolens* could only be partially sequenced but all key residues were be identified (63). Of note, one sequence from one individual (whom the initial cell line originated from) was determined for each species. Polymorphisms, as described for Rhinolophus, were not considered in this study (64). Multiple ACE-2 polymorphisms were indeed observed only in *Rhinolophus euryale* and only the major amino-acid is shown in the table.

## Acknowledgements

The authors acknowledge Erik Boehm for help with editing the manuscript and Pascal Cherpillod for facilitating the work in the BSL-3 laboratory at Geneva University Hospitals. We thank Joanna Damas and Harris A Lewin from the University of California for kindly providing the amino-acid sequence of *Pipistrellus pipistrellus* species (obtained from an original sequence from the Zoonomia Consortium). A549-ACE2 and VeroE6/TMPRSS2 cell lines were kindly provided by Pr Mirco Schmolke and Dr Beryl Sanchez at the University of Geneva.

## Author Bio

Dr Manel Essaidi-Laziosi is a Research and Teaching Fellow at the Faculty of Medicine of Geneva, Department of Microbiology and Molecular Medicine. Her main research interests are the study of the interactions of respiratory viruses with host cells, including differentiated human airway epithelia, during *in-vitro* and *ex-vivo* viral infections.

---

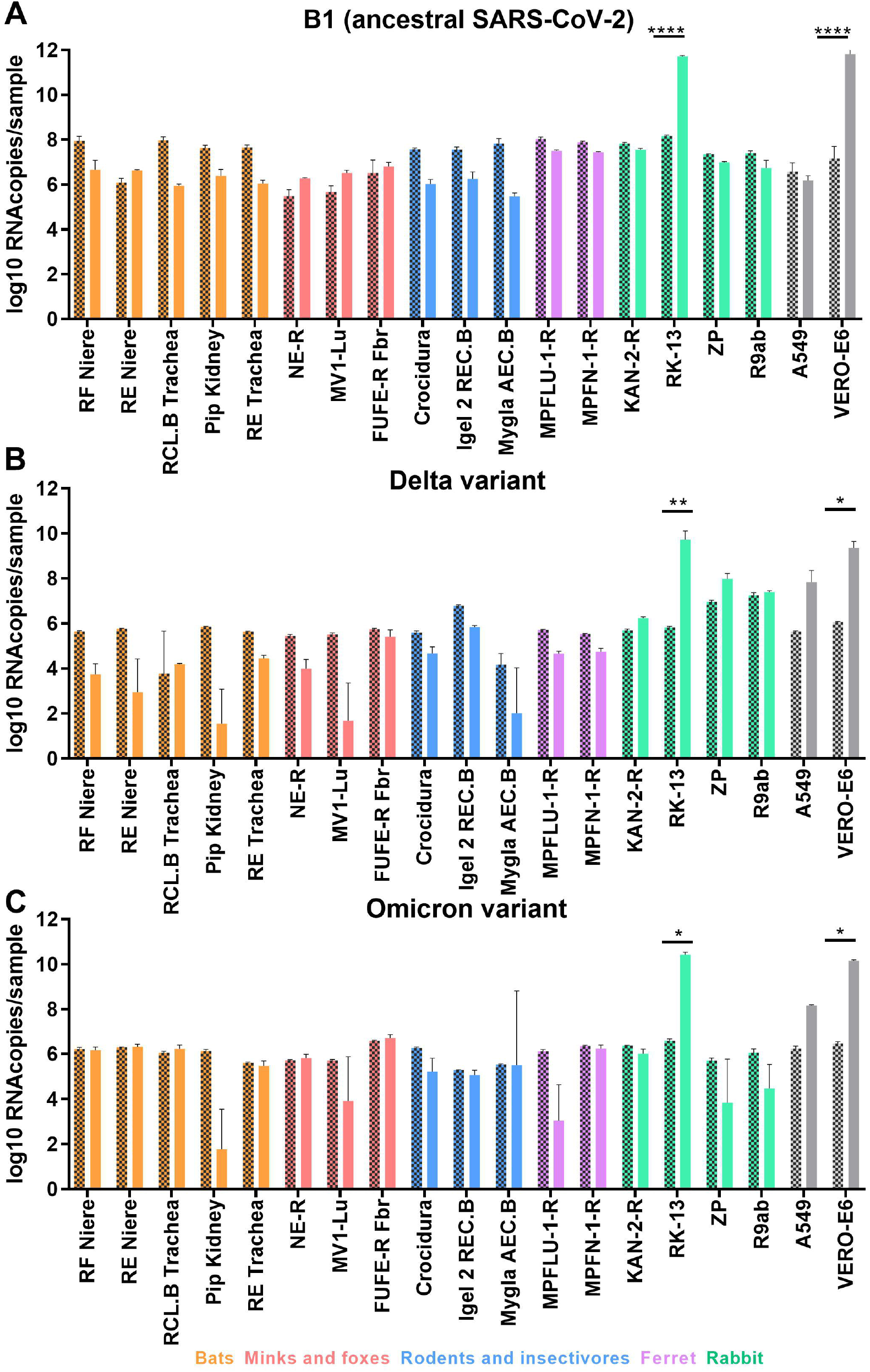

