## Supplementary material for "Distinct phenotype of SARS-CoV-2 Omicron BA.1 in human primary cells but no increased host range in cell lines of putative mammalian reservoir species": supp data

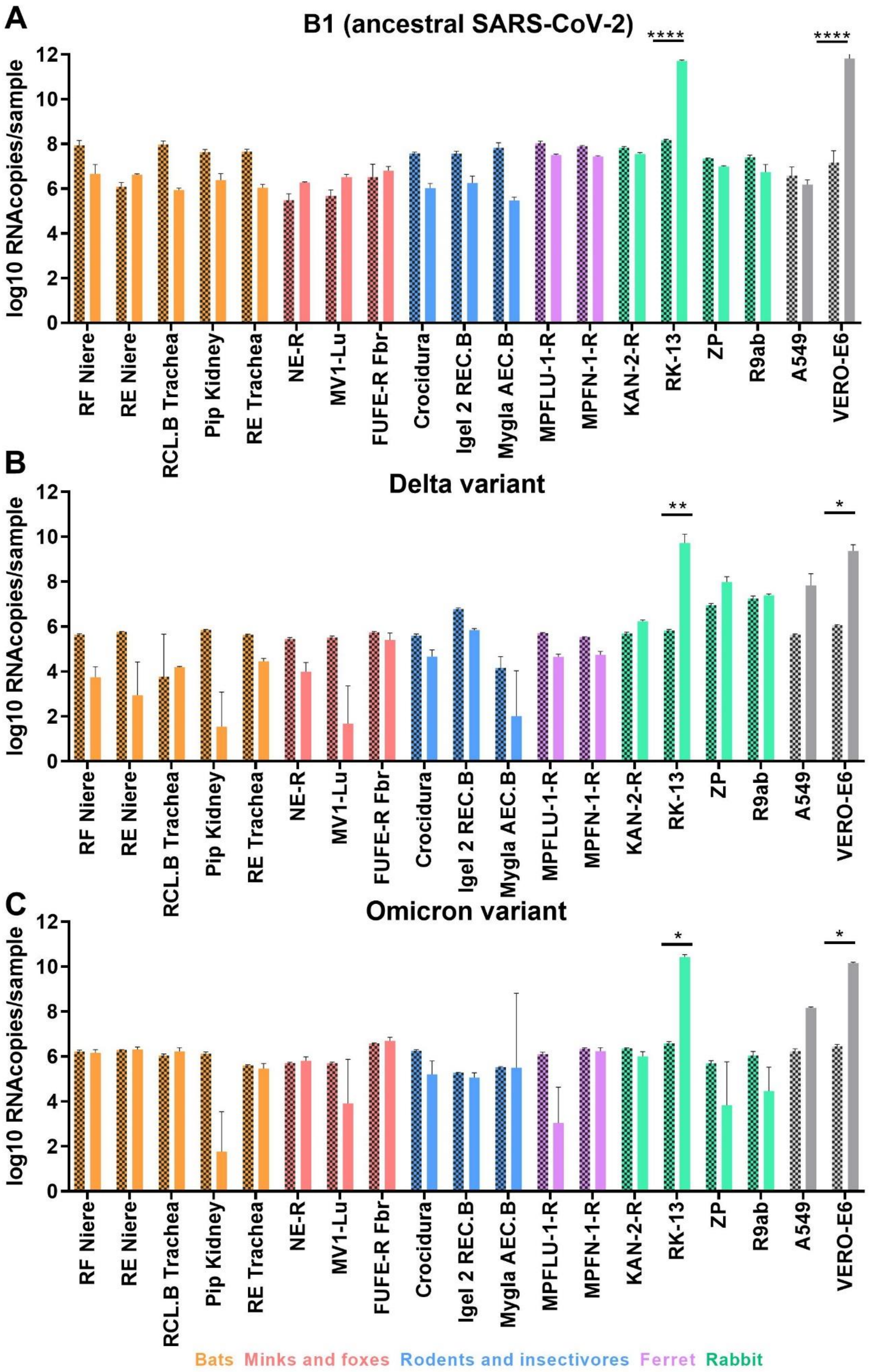
A
B1 (ancestral SARS-CoV-2)
B
Delta variant
C
Omicron variant
log10 RNAcopies/sample
****
**
*
RF Niere
RE Niere
RCL.B Trachea
Pip Kidney
RE Trachea
NE-R
MV1-Lu
FUFE-R Fbr
Crocidura
Igel 2 REC.B
Mygla AEC.B
MPFLU-1-R
MPFN-1-R
KAN-2-R
RK-13
ZP
R9ab
A549
VERO-E6
Bats Minks and foxes Rodents and insectivores Ferret Rabbit

**Figure S1: B1, Delta and Omicron SARS-CoV-2 variants replicate only in rabbit RK-13 cells.** From the same infections as in figure 3 of mammalian cell using clinical isolates of SARS-CoV-2 including the ancestral B1 lineage (A), Delta (B) and Omicron (C) variants, subgenomic viral RNA was quantified by RT-PCR from intracellular RNA at 1Hpi (hatched bar) to 96Hpi (open bar).. Statistical significance increase was calculated using 2-way ANOVA for fold change. *P < 0.05 and ****P < 0.0001 (N=3-4).

| Variant | Pango Lineage | Collection date | Sequence of patient sample form which isolate was derived (Gisaid ID) | Reference virus isolate |
|---|---|---|---|---|
| Ancestral SARS-CoV-2 | B.1 | February 2020 | EPI_ISL_414019 | Bekliz et al., Ulrich et al. |
| Alpha | B.1.17 | December 2020 | EPI_ISL_2131446 | Bekliz et al., Ulrich et al. |
| Beta | B.1.351 | January 2021 | EPI_ISL_981782 | Bekliz et al., Ulrich et al. |
| Gamma | P.1 | January 2021 | EPI_ISL_981707 | Bekliz et al. |
| Zeta | P.2 | January 2021 | EPI_ISL_897700 | Bekliz et al. |
| Delta | AY.122 | April 2021 | EPI_ISL_1811202 | Bekliz et al. |
| Omicron | BA.1 | November 2021 | EPI_ISL_7605546 | Bekliz et al. |

**Table S1. List of all tested SARS-CoV-2 lineages.**

| **Species** | **Primer name** | **Forward or Reverse** | **5'-3' Sequence** |
|---|---|---|---|
| *Myodes glareolus* | Arvicolinae_FWD | Forward | ATGYCACGYTCCTYCTGGC |
| | Arvicolinae_REV | Reverse | CTAAAATGAAGTCTGAACATCATC |
| *Rhinolophus (clivosus and Euryale)* | Rhino_R_570 | Forward | CATAATGGTATCCTCTTGCC |
| | Rhino_F_583 | Reverse | AGATGGCAAGAGGATACC |
| | Rhino_F_839 | Forward | TTGGACAAATCTGTACCC |
| | Rhino_F_898 | Forward | TACTGATGCAATGTTGAACC |
| | Rhino_R_1659 | Reverse | CTCAACATCTGGTGCAGC |
| | Rhino_R_1788 | Reverse | GACTTCCTGTTCTGCTCC |
| | Rhino_F_1835 | Forward | GAACACTGACTGGAGTCC |
| *Crocidura suaveolens* | CroSua_FW1 | Forward | SYTYCTCAGCCTTGYTGC |
| | CroSua_FW2 | Forward | CTGGCTSYTYCTCAGCC |
| | CroSua_REV | Reverse | TCTCGYTTCATCTCCCACC |
| | CroSua_REV2* | Reverse | GATTTTGTCTTCTGAGAGTGC |

**Table S2. List of primer for ACE-2 amplification and sequencing.** *used only for sequencing
